# *EZTraits*: a Programmable Tool to Evaluate Multi-site Deterministic Traits

**DOI:** 10.1101/2021.10.18.464896

**Authors:** Matt Carland, Madhuchanda Bose, Biljana Novković, Haley Pedersen, Charles Manson, Shany Lahan, Alex Pavlenko, Puya G. Yazdi, Manfred G. Grabherr

**Affiliations:** SelfDecode.com, 1031 Ives Dairy Road Suite 228 - 1047, Miami FL 33179, United States

## Abstract

The vast majority of human traits, including many disease phenotypes, are affected by alleles at numerous genomic loci. With a continually increasing set of variants with published clinical disease or biomarker associations, an easy-to-use tool for non-programmers to rapidly screen VCF files for risk alleles is needed. We have developed *EZTraits* as a tool to quickly evaluate genotype data (e.g., from microarrays) against a set of rules defined by the user. These rules can be defined directly in the scripting language *Lua*, for genotype calls using variant ID (RS number) or chromosomal position. Alternatively, *EZTraits* can parse simple and intuitive text including concepts like ‘*any*’ or ‘*all*’. Thus, *EZTraits* is designed to support rapid genetic analysis and hypothesis-testing by researchers, regardless of programming experience or technical background. The software is implemented in C++ and compiles and runs on Linux and MacOS. The source code is available under the MIT license from https://github.com/selfdecode/rd-eztraits

## INTRODUCTION

Many common health disorders are highly polygenic. This means that calculating an individual’s aggregate genetic risk is often unfeasible without the use of complex *polygenic risk scores* (PRS) [1]. However, there are a number of health-relevant traits for which smaller subsets of variants can account for disproportionately large amounts of phenotypic variance. These mono- or oligogenic traits are therefore amenable to simpler analytical approaches.

One illustrative example is the *APOE* gene, in which a two-SNP haplotype may modulate an individual’s risk of late-onset Alzheimer’s disease by approximately 15x [2]. Another example is the ability to digest lactose into adulthood, which can be fully predicted on the basis of just six SNPs in the *MCM6* gene, among which a single heterozygous- or homozygous-derived genotype implies lactose tolerance [3]. Similarly, dietary tolerance to fructose can be predicted by the presence of a few different combinations of homozygous mutations in the *ALDOB* gene [4].

Furthermore, small numbers of variants may also be useful for characterizing individual variability within specific biological pathways. One example is the *COMT* gene, in which various four-SNP haplotypes have been associated with significant differences in the biological activity of the gene’s product enzyme [5, 6]. Even in the absence of a direct link to a clinical phenotype, such genetic markers may serve as a useful “jumping-off” point for further investigations into the etiological structure of clinically relevant phenotypes.

Here, we present *EZTraits*, a tool specifically designed for non-programmers. *EZTraits* is intended to assist with searching VCF files for the presence of mono- or oligogenic traits and return their trait associations to the user, based either on our library of variant-trait associations or new, user-added conditions and associations. Thus, *EZTraits* allows genomic researchers to quickly and easily analyze a wide variety of phenotypes of clinical and scientific interest, regardless of their level of programming ability.

## METHODS

*EZTraits* evaluates variant combinations by internally building and interpreting *Lua* scripts. The Lua programming language [7] was designed with ease-of-use in mind and has been widely adopted by non-programmers for computer game modding and writing plug-ins, making it a natural choice for use by researchers both with or without a coding background.

There are two ways for users to build analyses with *EZTraits*: (a) by writing or modifying a Lua “snippet,” which contains pre-made variables for supplying key genotype and phenotype information; or (b) by writing a plaintext rule set that provides genotype and phenotype information by using simple concepts such as *all* and *any*, which allows for a more intuitive and compact representation. This conversion feature allows users to easily write in rsID-trait associations to use with *EZTraits* without any “coding” at all.

### Using Scripts

Users can write Lua snippets directly by providing the appropriate genotype and phenotype information. For genotypes, SNPs can be referenced using either their rsID or chromosomal position (following the syntax ‘chr1:6658743’). Phenotype information is entered by modifying two return variables: the floating-point variable ‘*risk*’; and the string ‘*comment*’ — both of which can be manipulated directly in the Lua script snippet. These two variables allow the user to flexibly provide either quantitative or qualitative phenotype data (or both), depending on the trait being analyzed.

For example, the snippet:

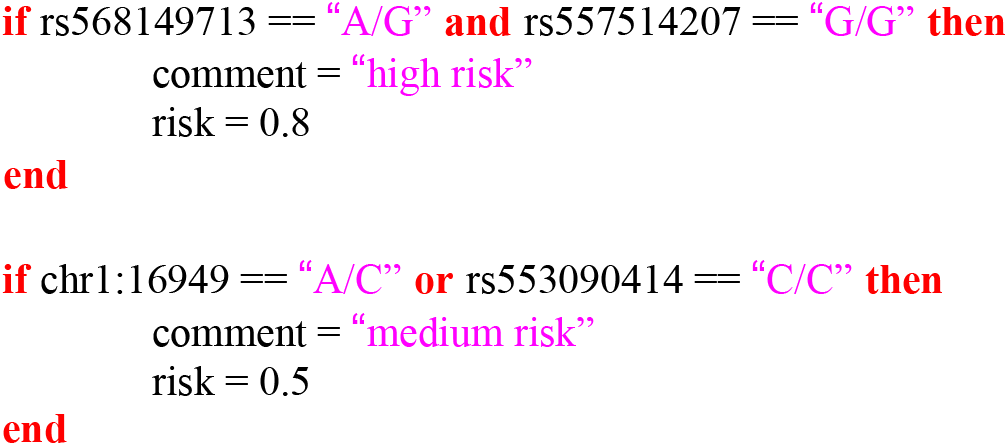

is completed into a valid Lua function by adding variables that correspond to the RS identifiers or chromosomal positions. These variables are automatically initialized from a VCF or TSV file, and together with a small amount of bracketing code, the complete function is:

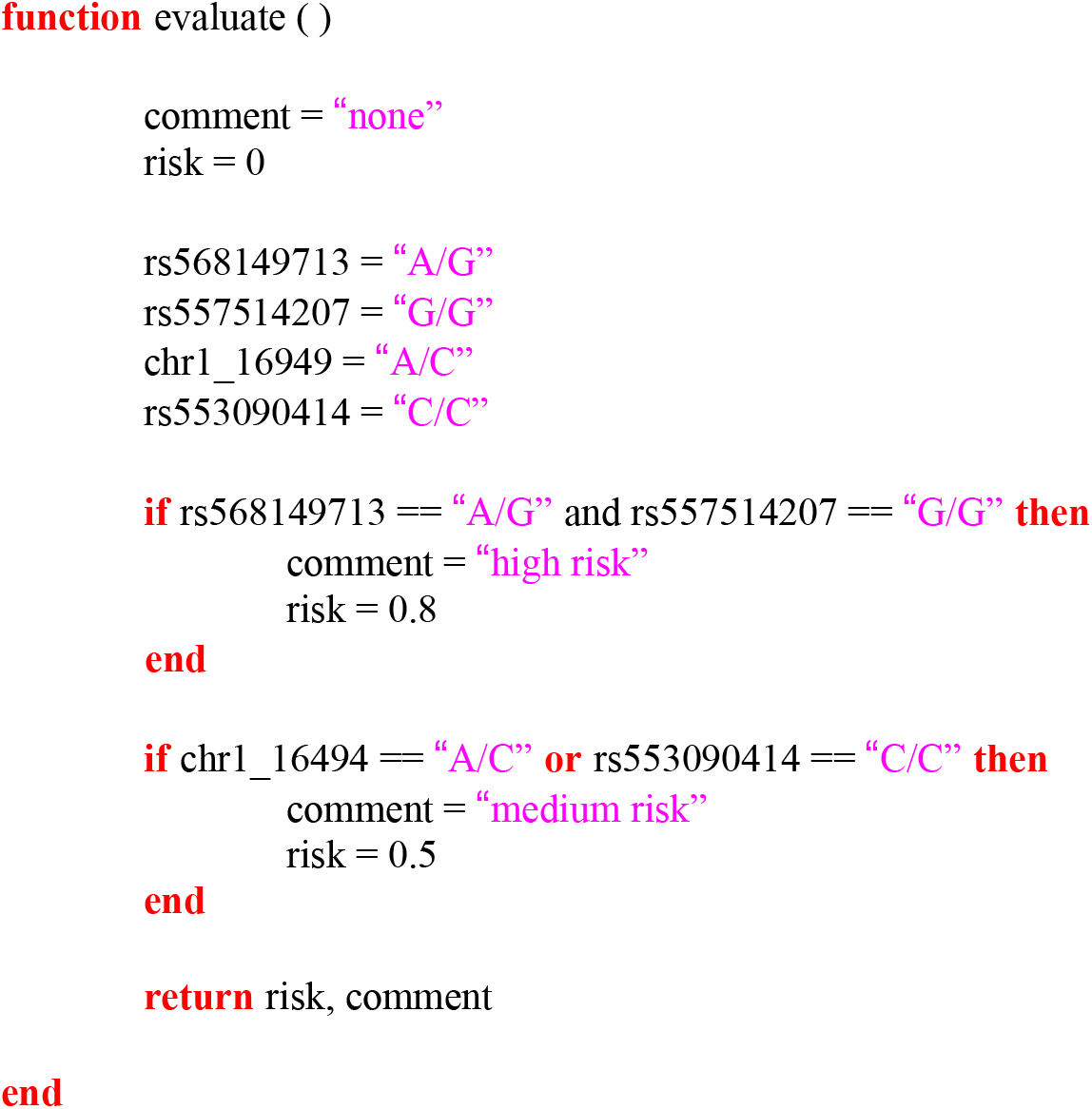

This function is then called directly from C++ by *EZTraits*, and the results are presented to the user.

### Structured text entry

In addition, *EZTraits* can automatically convert text files into Lua by applying some simple-yet-intuitive concepts, such as *any* and *all*, i.e., any of the following conditions satisfy a trait, or all in combination do. This text ruleset is then automatically converted into fully-functional Lua code via the tool *Txt2Lua*.

For example, the rules to define fructose tolerance/intolerance using three common causal SNPs can be written as:

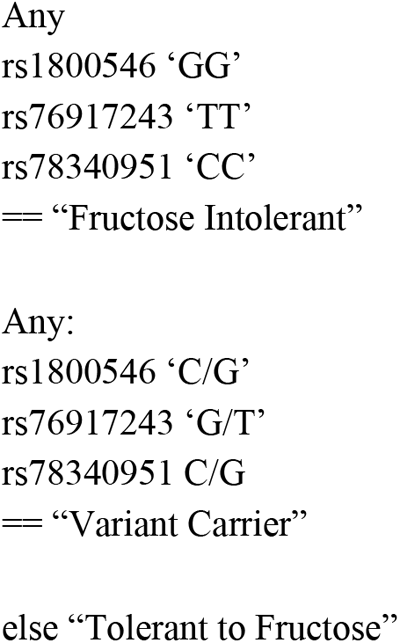

*EZTraits* accepts and interprets the keywords ‘All’, ‘Any’, and ‘else’, optionally followed by a colon. Acceptable genotype call formats include ‘CG’ and C/G (with optional single quotation marks), where the latter convention has to be used for sites that contain indels, e.g., ‘T/TGAT’.

The above text thus translates into the Lua snippet:

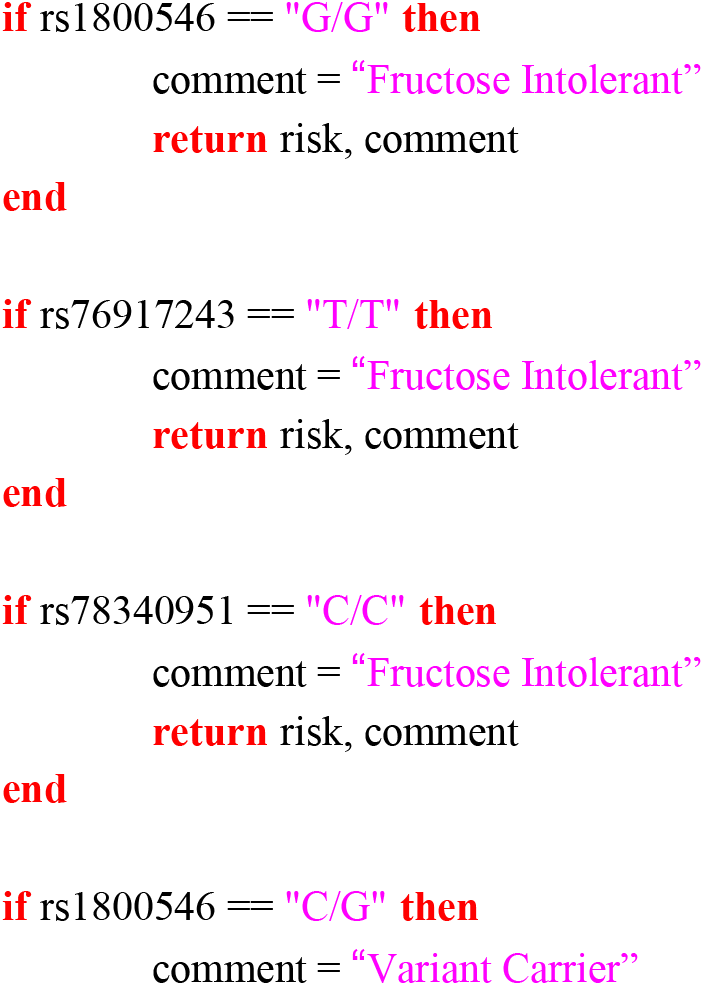

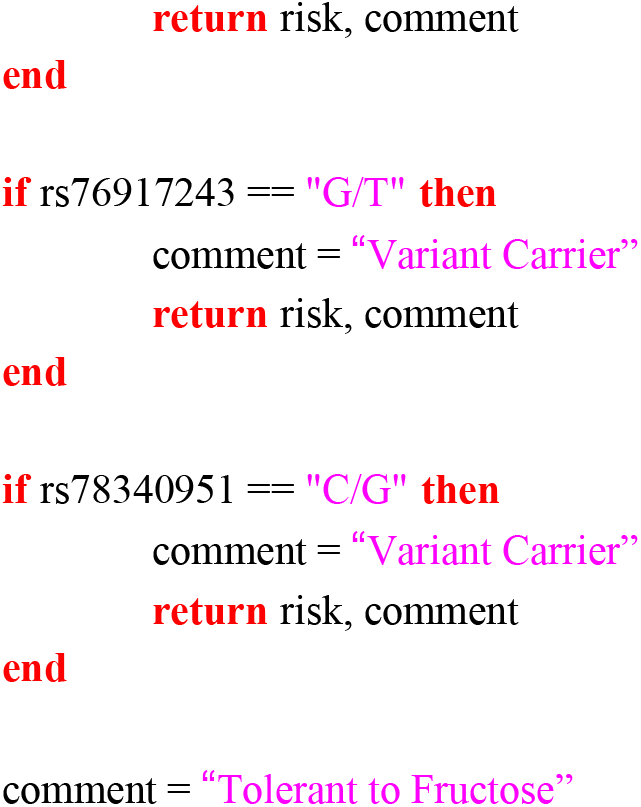

## RESULTS

*EZTraits* is a command-line tool that compiles and runs on Linux and Mac operating systems. Inputs are VCF or space/tab-delimited TSV files. The Lua interpreter, version 5.4.2, is embedded so that *EZTraits* has no external dependencies. *EZTraits* has minimal requirements in terms of RAM, using less than 5KB on average. It takes about 2.4 minutes to parse a whole-genome VCF file from the 1000 Genomes Project [8].

### Usage

*EZTraits* has two input parameters: (a) the VCF or TSV file; and (b) the Lua snippet. The usage for processing a VCF and TSV file is:

./EZTraits -i data/sample.vcf -lua scripts/test.lua
./EZTraitsCSV -i data/sample.csv -lua scripts/test.lua

The output is written to the console. To convert structured text, run e.g.:

./Txt2Lua -i scripts/fructose.txt > fruct_test.lua

